# The endoplasmic reticulum promotes microtubule organization and region-specific disassembly to execute Compartmentalized Cell Elimination

**DOI:** 10.1101/2025.05.08.652974

**Authors:** Karen Juanez, Hannah Selvarathinam, Madison Jones, Piya Ghose

**Affiliations:** The University of Texas at Arlington

**Keywords:** ER, MTOC, programmed cell death, neurodegeneration

## Abstract

Specialized cells, such as neurons, die during development and disease. How subcellular organization and interactions across diverse compartments direct death is not well-understood. We examine this by studying the Compartmentalized Cell Elimination (CCE) developmental death program of the *C. elegans* tail-scaffolding cell (TSC). We find endoplasmic reticulum (ER) shape genes and the microtubule (MT) severase SPAS-1/Spastin, all linked to neurodegeneration, cooperate to promote CCE. Super-resolution imaging reveals profound spatiotemporal dynamics of MTs and the ER across CCE, including enrichment in a stereotyped degenerative node. We observe an ER-dependent non-centrosomal microtubule organizing center (ncMTOC) in the degenerative node, where the ER locally promotes both MT organization and SPAS-1/Spastin-mediated MT severing. The ER is spatially confined, such that SPAS-1/Spastin also has an ER-independent role. Our study expands our understanding of the cell biology of specialized cell death during development and presents molecular links to pathological neurodegeneration, paving the way to neurodegenerative therapies.

## Introduction

Polarized cells, such as neurons, are defined by distinct compartments that can differ greatly in their internal architecture and their relative contributions to cell function(1). Specialized cells can have elaborate architecture and, like many cell types, are subject to programmed elimination, an important feature of normal development and homeostasis. For example, a large proportion of brain neurons are eliminated during development to refine and sculpt the brain (2). Specialized cells can also degenerate under pathological or injury states, which often share common morphological features with programmed cell regression and likely common molecular machinery(3). Failure in programmed neuronal regression can cause neurodevelopmental and psychiatric disease (4, 5). Programmed neuronal death and localized neuronal elimination (pruning) show morphological commonalities with neurodegeneration (6). Much of our growing knowledge of the molecular mechanism behind the various forms of pruning stems from *in vivo* studies in model organisms such as *Drosophila*(2, 7, 8) and *C. elegans*(9). The molecular mechanism for complex cell elimination broadly is, however, not fully understood. For example, gaps in our knowledge remain in understanding how the different subcellular domains of complex cells are differentially eliminated and the contributions of subcellular structures within them.

A particularly interesting and underexplored question pertaining to the mechanisms underlying developmental death of morphologically complex cells is the contribution of specific subcellular structures and their associations. There is growing interest in inter-organellar interactions and how these influence cell fate and function(10) and the impact of spatial distribution. How intracellular interactions among subcellular networks, such as the endoplasmic reticulum (ER) and the cytoskeleton, that occupy large, and diverse, expanses within cells, regulate cell death is not understood. Moreover, the spatiotemporal dynamics of these subcellular interactions in different compartments during the course of a complex cell elimination paradigm *in vivo* is not well-studied.

Among widely pervasive subcellular structures is the ER, the major intracellular membrane system formed of sheets and tubules that occupies the largest surface area in the cell. The ER performs a variety of functions in the cell and, for morphologically complex neurons, has been described as a “neuron within a neuron”(11), with smooth tubular ER that are very narrow making up axons and dendrites. The neuronal ER extends across the entire cell, including the extremities of the axons and dendrites(12). The ER’s intricate and elaborate structure is dependent on various ER-shaping proteins such as the hairpin proteins Lunapark and the dynamin-like GTPase Atlastin(12). The latter has been shown to be important for dendrite branching and function in *C. elegans*(13). Structural changes (from sheets to tubules) in the ER are known to take place during mammalian cell division(14). At the Drosophila neuromuscular junction, Atlastin has been shown to be important for synaptic function cell non-autonomously in the muscle(15).

Microtubules (MTs) are a key part of the cytoskeleton that are polymers of tubulin and an especially interesting structural component for the specialized cell, in which each cell compartment has a different structural demand according to function. During pruning, the local cytoskeleton is known to be disassembled, and it has been suggested that local organization of MTs and their disassembly serves as a local trigger to instruct where pruning occurs(16, 17). MT depolymerization has been associated with axonal beading/swelling and fragmentation in neurodegenerative disease(18). Interestingly, retraction bulbs and a failure of axon degeneration are observed when MTs are disorganized following pharmacological disruption in DRG cultured neurons(19). Several studies have described connections between the ER and components of the cytoskeleton. The ER has been implicated in theory in the spatial distribution of MTs(20) as a function of ER tubule junction dynamics to direct the distribution of MT bundles.

One feature of MTs is their ability to organize into specialized sites, called microtubule organizing centers (MTOCs), the best described being the centrosome(21, 22). Beyond the centrosome, “non-centrosomal microtubule organizing centers” are seen in differentiated cells performing functions such as axon regeneration(22) and there is increasing interest in what additional roles MTOCs play. Importantly, a role for ncMTOCs in programmed cell death has yet to be described. Various components of MTOCs have been identified(21, 22) in *C*.*elegans*. Outstanding questions include how MTs, which are held in place as arrays at their minus ends, are directed to forming MTOC and what factors stabilize this structure and anchor MTs(23). Work has shown that this may be through other MTs, such as the pre-existing MTs of the Golgi Apparatus(24). Open questions also remain as to a direct involvement of ncMTOCs in cell death, and how MTOCs are generated in specialized cells.

Another characteristic of MTs is that they undergo dynamic changes of severing and stabilization(25). MT severases(26) have been identified and have also been linked to neurodegenerative diseases such as Hereditary Spastic Paraplegia (HSP)(27). A MT severase that has gained particular attention is the ATPase Spastin(25, 28, 29), given its recent additional implications in the opposing function of MT stabilization(30). *In vivo* physiological contexts demonstrating this dichotomy are not well described nor what regulates Spastin’s dual functional capabilities. Spastin has been shown to function *in vivo* in synapse regulation(31, 32). Spastin has also been associated with the endoplasmic reticulum (ER) (18). Studies have shown that Spastin physically interacts with Atlastin(33), thus also linking the ER to MT regulation.

We have previously described a non-apoptotic specialized cell death program, Compartmentalized Cell Elimination (CCE) in the nematode *C. elegans(34)*. We have observed this highly organized stereotyped program in two polarized cell types in the nematode embryo, the tail-scaffolding cell (TSC) and the CEM sensory neurons. CCE is characterized by distinct elimination morphologies in the different compartments in the same cell and bears morphological features resembling developmental pruning and neurodegeneration.

Here we structurally and functionally characterize a cell compartment-specific degenerative hub in the dying TSC where the ER plays dual roles in MT organization and disassembly important for cell dismantling. We show roles for conserved genes known to be linked to neurodegenerative disease to play important roles in developmental cell death. We highlight the importance for spatiotemporally regulated sub-cellular interactions and present the ER as a killer organelle with dual roles, MT organization and MT dismantling. We show that the ER is spatially confined, and suggest that MTs are regulated by Spastin in a compartment-specific, ER-dependent and ER-independent manner. Finally, we present the ER in a previously undescribed role in ncMTOC formation. Our work helps bridge gaps in our knowledge about specialized cell elimination and subcellular events and suggests molecular links between developmental neuron death and cell death during neurodegenerative disease setting the stage for the identification of novel therapeutic strategies.

## Results

### Defining the degenerative hallmarks of Compartmentalized Cell Elimination (CCE)

We have previously described Compartmentalized Cell Elimination (CCE) as an ordered, organized killing program. To probe deeper, we evaluated the distinctive compartments of the TSC during TSC development and during CCE in greater detail. (**FIG 1A)** Shows schematic of TSC compartments and key CCE steps. We imaged our previously-described membrane marker for the TSC (TSC-myrGFP)(34). The TSC has the following compartments distinguishable as CCE progresses: the soma (s), the soma-process junction (spj), the proximal process segment (p), and the distal process (d), comprising the distal node (dN), the distal process tip (dT). We define five morphologically distinct stages of TSC development and death (**FIG 1B-G**). The “IY” (intact, young) cell (**FIG 1B**) extends its single process posteriorly forming a sharp spike that scaffolds the tail tip. The TSC matures to the “IMA” (intact, mature) stage, now with an elongated process that is uniform, without a distinct spike (**FIG 1C**). The first notable sign of CCE is an initial nicking event at the soma-process junction (spj) (**FIG 1C**). Following this, at the “BA” (beading, attached) stage, the cell can be seen with three disparate compartments. The cell body (s) appears round superficially resembling an apoptotic corpse early on. The proximal process (p) undergoes beading, still loosely attached to the soma (**FIG 1D**). The beaded proximal process (p) then fragments, and the process and soma become clearly disconnected in the “BD” (beading, detached) stage (**FIG 1E**). The proximal segment beading and fragmentation is reminiscent axon degeneration following injury(18). Starting from the beading stages, distally, a hallmark of CCE is the formation of a varicosity, a distal node (dN), that we identify as a stereotyped degenerative hub **(FIG 1D**). As CCE progresses, the proximal process segment beads are removed fully first. The distal process tip (dT) appears to retract into the distal node (dN) at the “SDR” (soma-distal retracting) stage (**FIG 1F)** in a manner morphologically similar to axonal retraction following actin disruption(18). Ultimately, the soma and distal remnant are stochastically removed by phagocytosis by neighboring cells at the SDD (soma-distal degrading) stage (**FIG 1F**). The distal node remains uncharacterized, and we posit that defined subcellular events important for CCE take place here.

**Figure 1.**
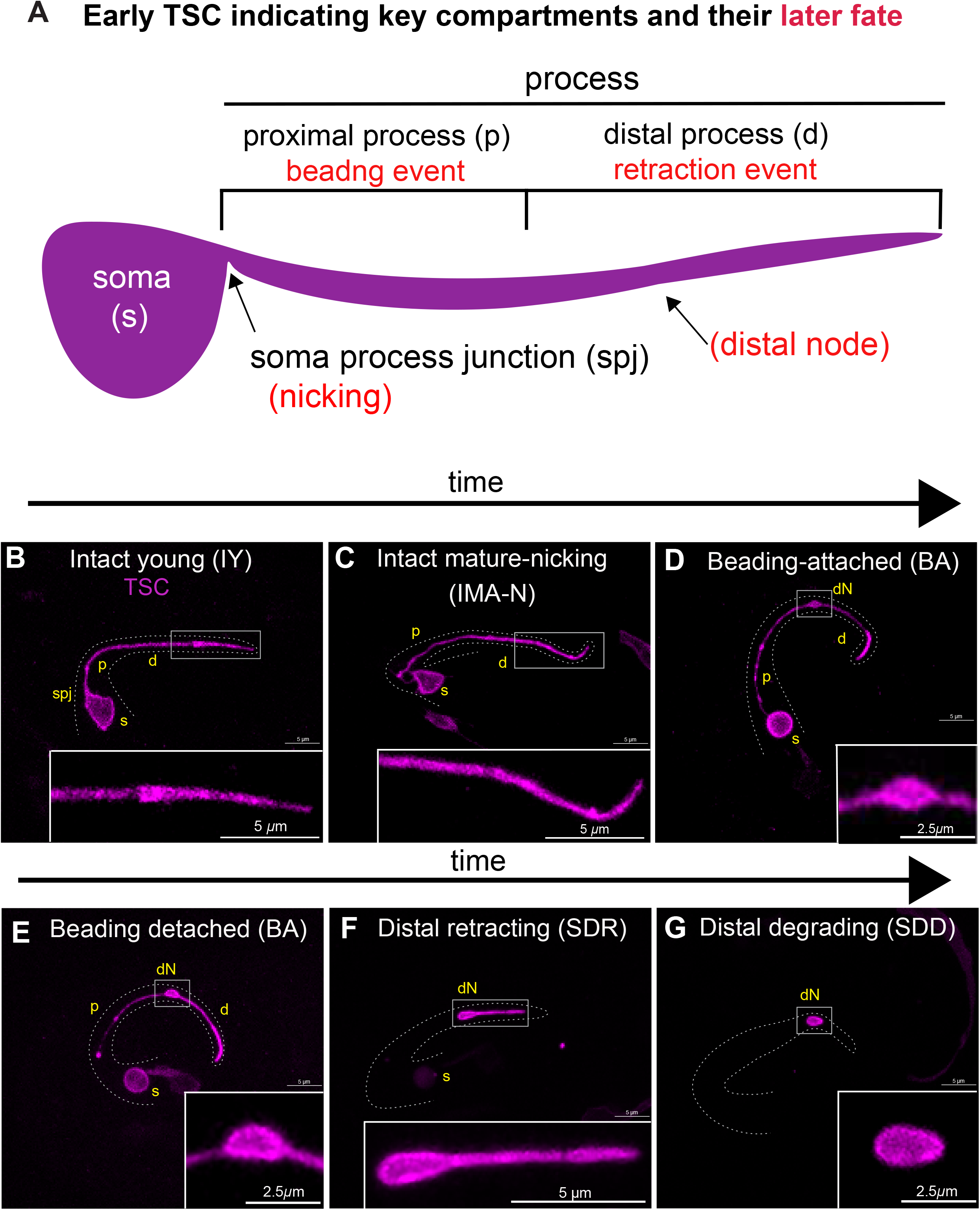
General account of TSC compartments and CCE stages. **(A)** Map of TSC indicating key regions of events as seen in young intact. Red, later events of CCE. **(B-G)** CCE stages: (**B**) young intact (IY) ∼600 mpf, (**C**) mature intact to nicking (IMA-N) ∼650 mpf, (**D**) beading-attached (BA) 655 mpf, (**E**) beading detached (BD) ∼670mpf, (**F**) soma distal retracting (SDR) ∼ 745 mpf and (**G**) soma distal degrading (SDD) ∼800 mpf. s, soma; p, proximal process segment; spj, soma process junction; d, distal process segment; dN, distal node; dT, distal tip.

### ER shape genes and the MT severase SPAS-1/Spastin promote CCE

To understand CCE at the molecular level, we performed a forward genetic screen of 27,000 F2 worms using a TSC membrane reporter GFP transgenic strain used in our earlier studies (34). From this screen, we obtained a mutant, *ns824*, **(FIG 2A)** which has an array of CCE defects, predominantly a persisting distal node (**FIG 2C**). From Whole Genome Sequencing we found a C is mutated to T change resulting in P332 changing to S in exon of the gene *lnp-1* (**Supplemental FIG 1A**), which encodes the homolog of mammalian Lunapark, a hairpin membrane fusion protein in the endoplasmic reticulum (ER) membrane that aids the stabilization of ER three-way junctions and the maintenance of ER shape(35). We tested another *lnp-1* allele, *lnp-1(tm1247)*, and found this to phenocopy (**FIG 2B**). We also found *lnp-1* to be expressed in the TSC using a promoter fusion construct (**FIG 2D, D’**) and functions cell autonomously in the TSC from cell-specific rescue experiments (**FIG 2E**). We proceeded to test other ER shaping genes(13). We tested the dynamin-like GTPase ATLN-1/Atlastin. Mammalian Atlastin is thought to function with Lunapark to stabilize ER three-way junctions and is needed for ER tubule formation. We found that *atln-1(ok1144)* had a range of CCE defects similar to those seen in *lnp-1* mutants (**FIG 2F**). Here too we found that *atln-1* is expressed in the TSC and functions there (**FIG 2H, H’)**.

**Figure 2.**
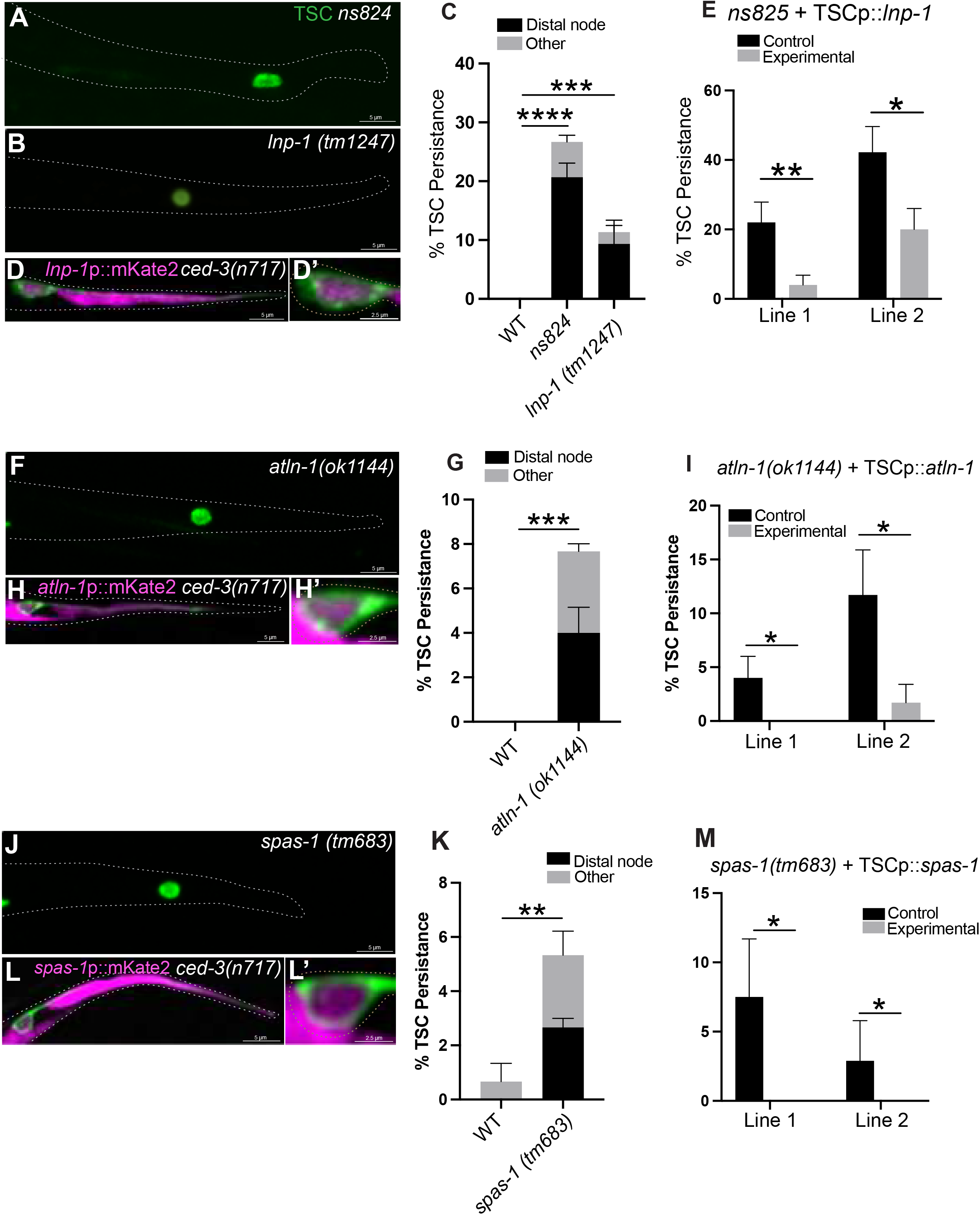
ER shaping genes and the microtubule severase SPAS-1/Spastin promote CCE. **(A)** Distal node (dN) phenotype of *lnp-1(ns834)* mutant obtained from CCE screen **(B)** Distal node (dN) phenotype of *lnp-1(tm1247)* allele. **(C)** Quantification of *lnp-1* mutant phenotypes **(D**,**D’)** Expression of *lnp-1*p::mKate2 in the TSC. **(E)** TSC Cell -specific rescue of *lnp-1(ns834)*. **(F)** *atln-1(ok1144)* mutant <u>distal node (dN)</u> phenotype. **(G)** Quantification of *atln-1* mutant phenotypes. **(H)** Expression of *atln-1p*::mKate2 in the TSC. **(I)** TSC Cell - specific rescue of *atln-1(ok1144)*. **(J)** *spas-1* mutant phenotype. **(K)** Quantification of *spas-1* mutant phenotypes. **(L)** Expression of *spas-1p*::mKate2 in the TSC. (**M**) TSC Cell -specific rescue of *spas-1(tm683)*. N>50. ns (not significant) *p* > 0.05, * *p* ≤ 0.05, ** *p* ≤ 0.01, *** *p* ≤ 0.001, **** *p* ≤ 0.0001.

We next probed the molecular link between ER shape and CCE. Previous studies report that mammalian Atlastin binds to Spastin(33), a dual-functioning ATPase(30) that has been shown, classically, to sever, but more recently reported to also stabilize MTs. We looked at a mutant for *spas-1* which encodes the *C. elegans* homolog of Spastin, SPAS-1. We found that *spas-1* mutants also had similar CCE defects (**FIG 2J,K**) that can be rescued TSC-cell-specifically **(FIG 2M)**. *spas-1* is also expressed in the TSC ((**FIG 2L, L’)**. From epistasis studies, we found that, in terms of total numbers, the mutant combinations did not have additive effects **(Supplementary FIG 1B)** suggesting a common pathway. We do note that the penetrance of the defects in all three mutants is low, which has been reported (personal communications) which may be due to the hypomorphic nature of the viable alleles and other compensatory effects. Having identified regulators of the ER and MTs as new and important molecular components of CCE, we proceeded to employ super-resolution microscopy to cell biologically and mechanistically characterize how the ER and MTs may interact to promote CCE.

### CCE involves organized microtubule dynamics regulated by ER shape genes and SPAS-1/Spastin

We generated a MT reporter for the TSC by tagging GFP with TBA-1 (alpha tubulin) and examined it against a plasma membrane marker (myr::mCherry) in wild-type embryos across CCE stages. We found that prior to CCE, the TBA-1 signal of intact young (IY) appears to be enriched in the distal spike (**FIG 3A**). Just prior to death onset, TBA-1 appears to be more discretely localized. At the soma-process “severing/nicking” step (**FIG 3B**), proximally, the TBA-1 signal became highly localized (movement toward the node region). As this proximal segment starts to bead, TBA-1 is enriched at the beads (**FIG 3C**). Distally, TBA-1 was highly enriched in the distal node (**FIG 3C**) from this stage with signal accumulation intensifying as CCE progresses. Given the higher trend of distal node persistence of the ER shape mutants, and our poor understanding of neurite retraction as a pruning mechanism, we directed our focus on the events of distal process elimination and the role of the distal degenerative node.

**Figure 3.**
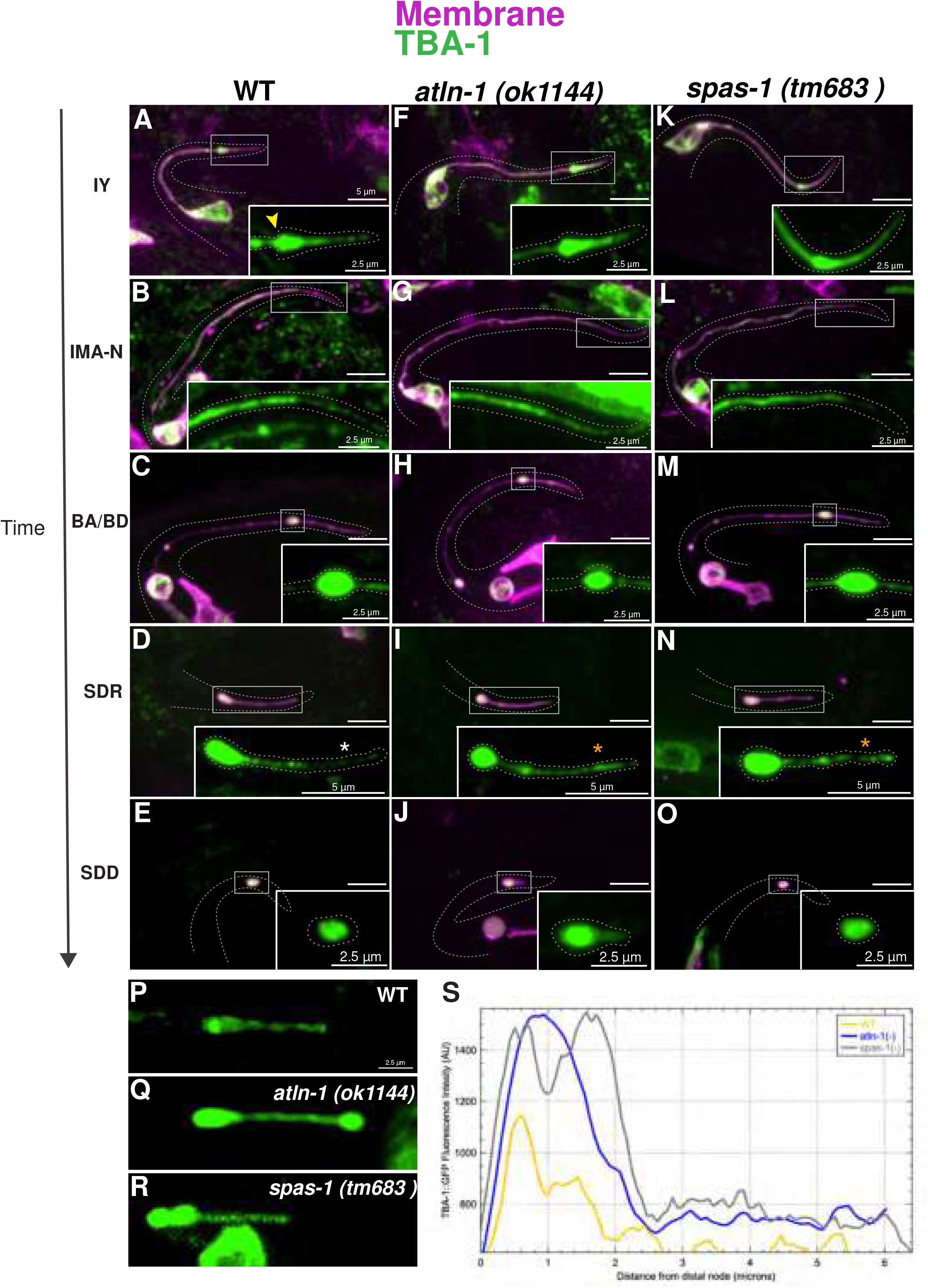
Microtubule dynamic changes during CCE depend on ATLN-1/Atlastin and SPAS-1/Spastin. TSC full cell with distal process magnified in inset. TSC TBA-1 dynamics across CCE stages in wild-type **(A-E)**, *atln-1* mutants **(F-J)** and *spas-1* mutants **(K-O)**. Accumulation of TBA-1 in intact young (IY) spike shown by yellow arrowhead **(A)**. TBA-1 signal appears less in soma distal retraction (SDR) distal tip (dT) WT than in *atln*-1**(I)** and *spas*-1 mutants and **(N)** as shown by white and orange arrowhead. Distribution of TBA-1 in wild-type, *atln-1 (ok1144)* and *spas-1(tm683)* mutants **(P-S)** Quantification of TBA-1 distribution of via intensity plot profile. Scale bar: 5 or 2.5 μm, Images are representatives, N>5.

We then tested TBA-1 in *atln-1(ok1144)* and *spas-1(tm683)* mutants (**FIG 3F-O**). We saw clear differences from IMA-N to SDD in *atln-1* (ok1144) **(FIG3H-J)** and *spas*-*1* (tm683) (**FIG 3M-O**). While TBA-1 is enriched in the distal node in all genotypes (**FIG 3C-O**), the intensity in the distal node (dN) and distal tip (dT) was higher in mutants. We quantified this difference (**FIG 3P-S**). CCE also appears to arrest at the “SDR” stage of CCE when MTs are stabilized, with the distal process failing to retract fully and resolve into the distal node (**FIG 2**). This suggests MT dismantling facilitated by ER shape genes and SPAS-1/Spastin is important for distal process retraction.

### The ER is spatially-restricted and has stereotyped dynamics during CCE

Given the importance of ER shape genes for MTs during the distal retraction stage of CCE, we proceeded to look at the localization of the TSC ER in wild-type embryos across CCE stages (**FIG 4A-E**). We visualized the ER using ER vit22(KDEL::oxGFP) (template: a gift from Barth grant) (**FIG 4A-E**). The ER appears to be uniform in the TSC process early on, which we speculate to represent ER tubules. Curiously, the ER does not extend to the distal tip of the process at any stage suggesting that there are ER-independent events at the distal tip (FIG A-E). The ER is enriched at the distal node during the “beading” stages (**FIG 4E**) and is tightly restricted in this node, never extending further distally. We proceeded to look at the ER in *atln-1* mutants. Loss of Atlastin is known to cause aberrant ER structure(13). While ER in Atlastin mutant shows aberrant structure **(FIG 4F-J)**, the ER remains confined and absent of the distal tip (**FIG 4F-J**). Looking at the distal node specifically, we notice more stronger signal in wildtype (**FIG 4C**) than in *atln-1* mutants (**FIG 4H**). This suggest that Atlastin’s role in ER stability may be important for CCE events in the distal node.

**Figure 4.**
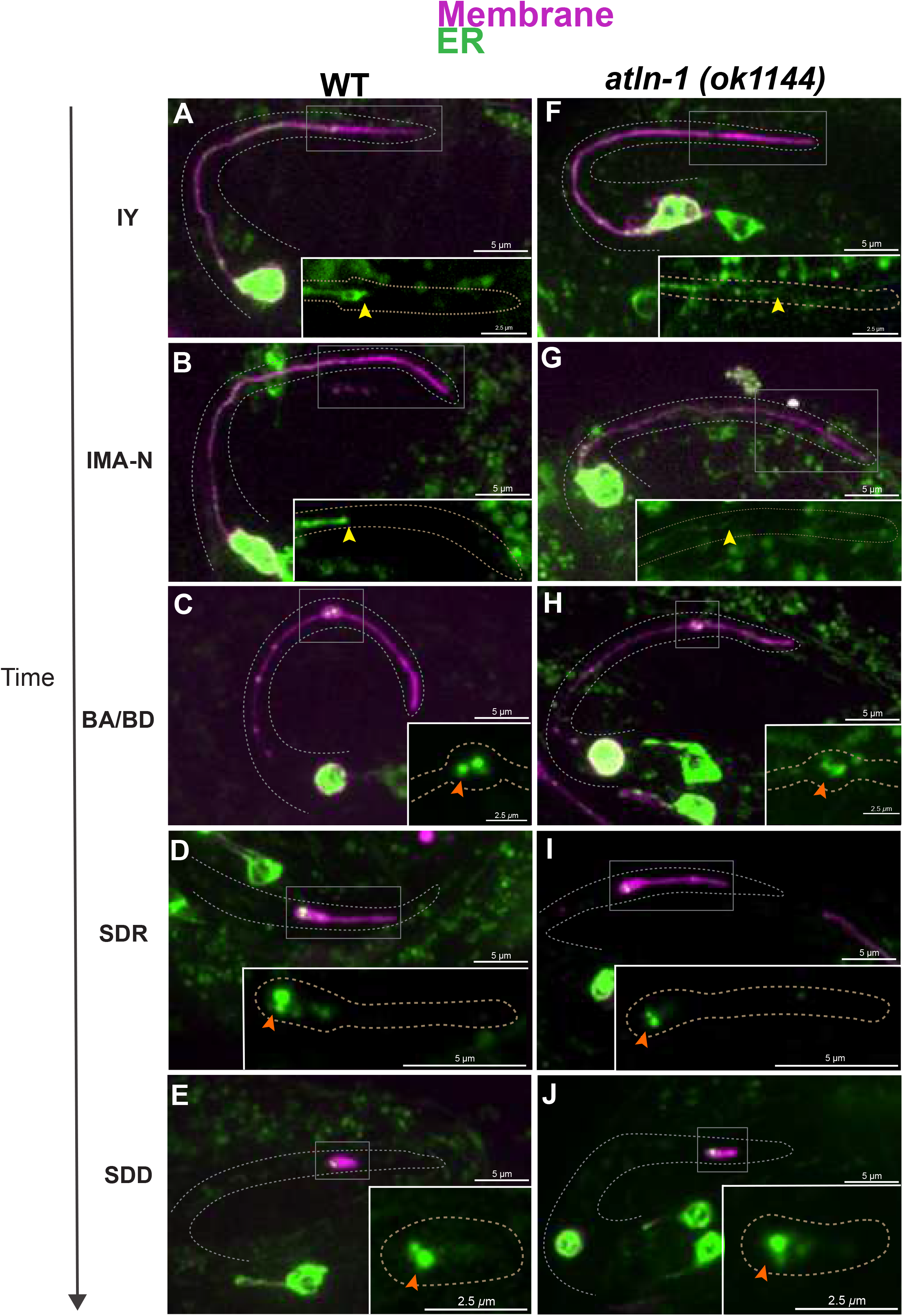
ER morphology in the TSC is altered in the absence of ATLN-1/Atlastin. TSC ER(KDEL) dynamics across CCE stages in wild-type **(A-E)** and *atln-1* mutants **(F-J)**.Full cell and inset showing magnified distal process. ER appears restricted in distal tip (dT) shown by yellow arrow and localized in distal node (dN) once CCE progresses, orange arrow in both WT and *atln*-1 mutants. Scale bar: 5 or 2.5 μm, Images are representatives. N>5.

### The ER promotes ncMTOC formation in the distal node during CCE, but not during TSC process outgrowth

We next aimed to further define the subcellular nature of the distal node. Given the enrichment of MTs in the distal node as CCE progresses prior to its resolution, we asked whether a CCE-promoting non-centrosomal microtubule organizing center (ncMTOC)(21, 22) is present there. We hypothesized that this may be the ER itself, or regulated by it, given the ER is also enriched there. To test the hypothesis that the distal node harbors a bonafide ncMTOC, we employed the MTOC reporter GIP-1/GCP3/Gamma Turc (36) with endogenously tagged GFP and TSC cell-specifically GFP (**FIG 5A-G**). We looked at these markers in the TSC process during development and death. Interestingly, we found enrichment at two time points: earlier in TSC development prior to CCE onset (**FIG 5A**) and then again in the distal node of the dying cell (**FIG 5E**). This suggests that two temporally distinct ncMTOCs form in the TSC process, one during process outgrowth and one during process demise. Strikingly, while the MTOC marker enrichment of the pre-CCE TSC was not affected by loss of multiple ER shape genes, marker accumulation was lost in the distal node of the dying cell (**FIG 5F-G**). Notably, the ER is not localized in the early TSC process, but becomes discretely localized as CCE sets in, accumulating in the distal node (**FIG 4E**). The MTOC marker was notably less in the distal node in ER shape mutants *atln-1, lnp-1*, mutants (**FIG 5F-G**). This suggests the ER serves as an ncMTOC in the dying TSC only, not earlier and that early-stage MTOC is ER-independent, and that its stability and discrete localization is essential for this role. This, to our knowledge, is the first direct evidence of the ER’s involvement in ncMTOC formation as a function of its temporal localization. Thus, we observe that the ER plays both a constructive role in MT organization, as well as a destructive role in promoting SPAS-1-mediated MT severing, with both functions relying on ER stability and leading to the dismantling of the TSC distal node during CCE. Given the common enrichment of the ER, MTs and MTOCs markers and the involvement of ER shape genes we propose that the ER serves in a previously undescribed role in ncMTOC formation in this degenerative node. Of note, a bonafide ncMTOC is the Golgi (37), which also harbors Atlastin. To distinguish if the ncMTOC is ER or Golgi, we tested other ER shaping genes and mutants for the Golgi gene *aman-2* (which encodes mannosidase) for CCE defects. While the ER shaping genes did have CCE defects, the *aman-2* mutants did not (**FIG 5H**). This data supports that the ncMTOC is ER rather than Golgi.

**Figure 5.**
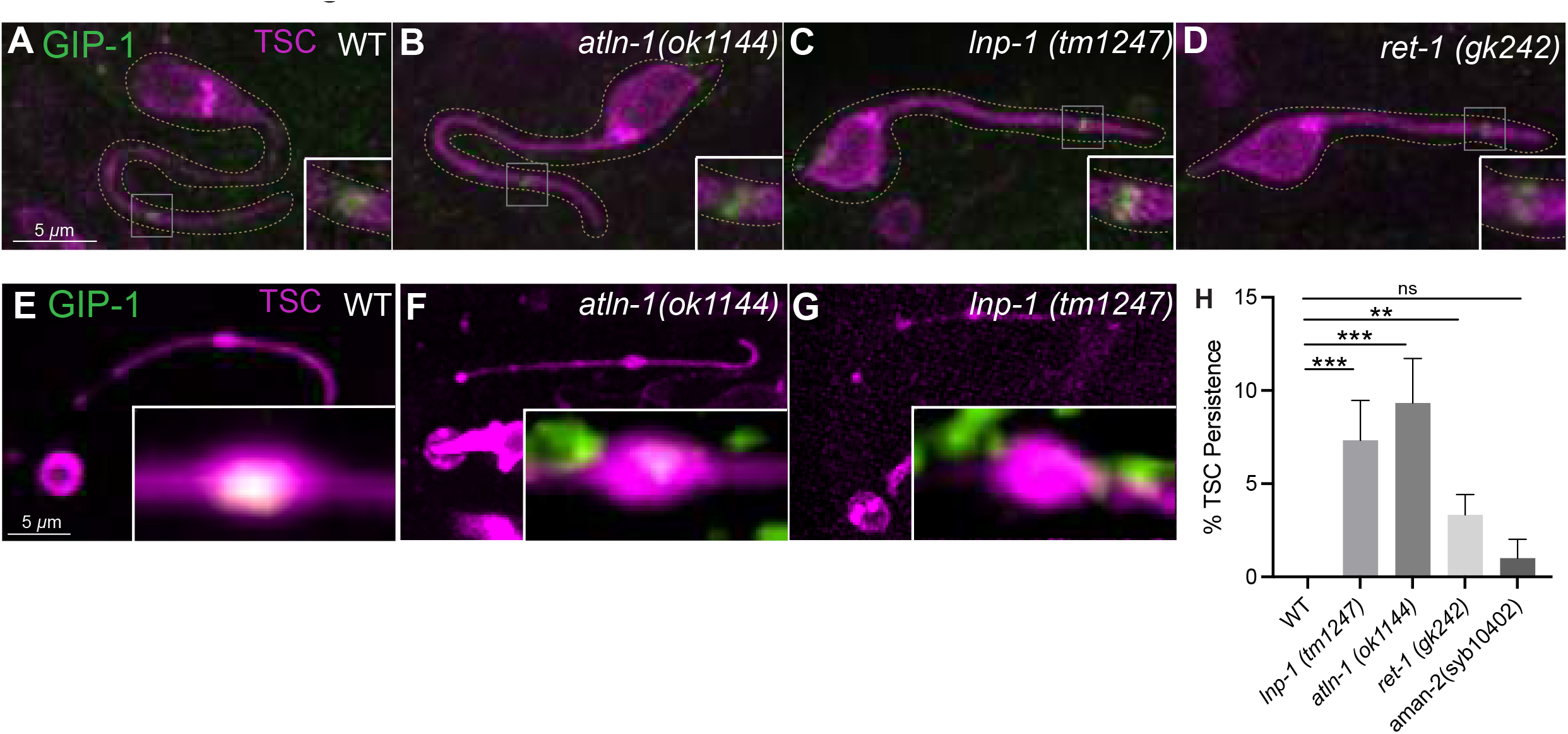
An MTOC is formed in the TSC distal node during CCE. GIP-1 localization in TSC (full cell and magnified in inset) at the intact young (IY stage) in (**A**) wildtype, ER shape gene mutants **(B)** *atln-1 (ok1144)*, **(C)** *lnp-1(tm1247)* and**(D)** *ret-1(gk242)* and at the beading attached stages in **(E)** wildtype **(F)** *atln-1 (ok1144)*, **(G)** *lnp-1(tm1247)*. **(H)** Quantification of ER genes VS Golgi mutant *aman-2* mutant. N>50. ns (not significant) *p* > 0.05, * *p* ≤ 0.05, ** *p* ≤ 0.01, *** *p* ≤ 0.001, **** *p* ≤ 0.0001. Scale bar: 5 or 2.5 μm, Images are representatives. N>5.

### ATLN-1/Atlastin regulates SPAS-1’s association with TBA-1 in distal node

Finally, to test the idea that ER integrity and organization and ATLN-1 direct SPAS-1 physically to cleave MTs in an organized fashion, we examined how ATLN-1/Atlastin regulated the engagement of SPAS-1/Spastin with TBA-1 We looked at SPAS-1 localization relative to TBA-1 in wild-type versus *atln-1(ok1144)* (**FIG 6 A-C, K-M**). In the distal node, in which the ER is enriched in wild-type but weaker in *atln-1* mutants, TBA-1 signal appears uniform in wild-type (**FIG 6D,E**), but appears disorganized and disproportionately distributed in *atln-1* mutants (**FIG 6N,O**). In the distal node, we observed various SPAS-1 localization differences. In wild-type, SPAS-1 appears to encompass TBA-1 but otherwise uniformly localized (**FIG 6F,G**). In compared to wild-type, SPAS-1, in *atln-1* appears to localize differently, more disorganized and disproportionate, appearing unevenly enriched at regions where TBA-1 is exaggerated (**FIG 6 P,Q**). Moreover, unlike wild-type (**FIG 6H-J**) TBA-1 appears to not be fully encompassed by SPAS-1 in *atln-1* mutants (**FIG 6R-T**). This highlights the organized nature of CCE MT dismantling. TBA-1 in the distal tip of the TSC process in wildtype appears more discreate (**Supplemental Figure S2)** but in the Atlastin mutant it appears more localized. SPAS-1 in this region appears uniform (**FIG 6B)** while less intense/absent in absence of ATLN-1(**FIG 6L**). This suggests that SPAS-1 is regulated in a manner dependent on ATLN-1, but independent of the ER in the distal tip, suggesting dual functions of SPAS-1 and a function that may be other than MT regulation.

**Figure 6.**
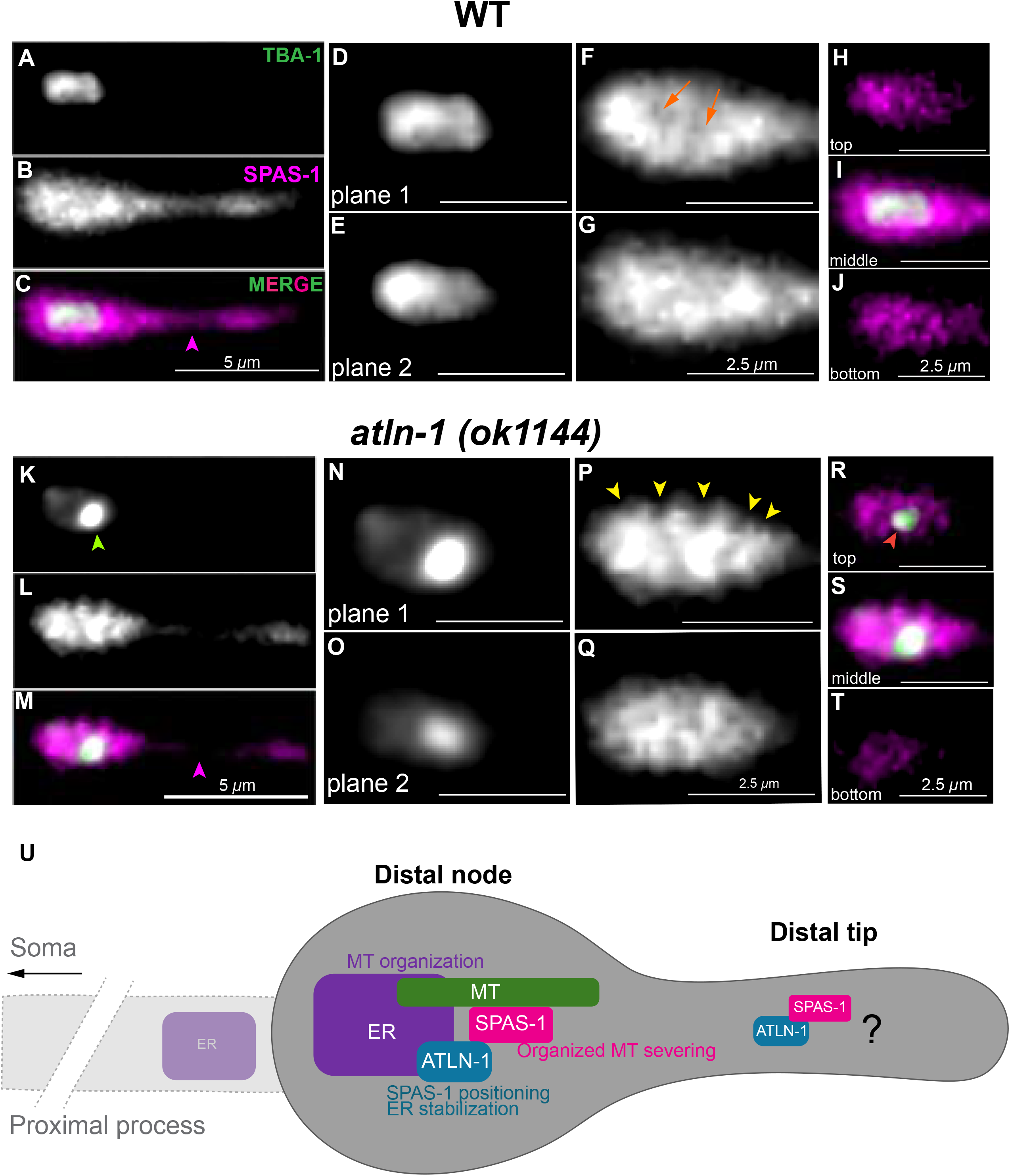
Organized association of SPAS-1/Spastin with TBA-1 depends on ATLN-1/Atlastin. TSC distal process at soma distal degrading (SDD) stage showing TBA-1 tagged with GFP and SPAS-1 tagged with mCherry. Wild-type: **(A)** TBA-1, **(B)** SPAS-1, **(C)** Merge.**(D,E)** two planes of TBA-1, **(F, G)** two planes of SPAS-1, **(H-J)** merge showing TBA-1 and SPAS-1 association suggesting SPAS-1 can encompass TBA-1. *atln-1 (ok1144)*: **(K)** TBA-1, **(L)** SPAS-1, **(M)** Merge. **(K)** <u>TBA-1 signal concentration indicated by green arrow.</u> **(N, O)** two planes of TBA-1, **(P, Q)** two planes of SPAS-1, **(R-T)** merge showing TBA-1 and SPAS-1 association. **(S)** exposed TBA-1 suggests SPAS-1 cannot properly engage with TBA-1 in *atln-1 (ok1144)* mutants. SPAS-1 appears broadly distributed across the TSC distal process in wild-type **(B)** but less visible in *atln-1 (ok1144)* **(L)** as shown by magenta arrowhead. TBA-1 appears more uniform in wild-type **(D, E)** and unevenly enriched in *atln-1 (ok1144)* **(N,O)**. SPAS-1 appears more uniform forming a cup in wild-type **(F, G)** and uneven with heavily accumulated areas near the TBA-1 enrichment *atln-1 (ok1144)*, shown by yellow arrowheads **(P,Q). (U)** Model: in the TSC distal node, the ER plays roles in both microtubule organization and disruption. Atlastin promotes ER stability and positions Spastin for organized microtubule severing. In the TSC distal tip Atlastin regulates Spastin in an ER independent manner in a noncanonical function of Spastin, independent of microtubules. Scale bar: 5 or 2.5 μm, Images are representatives. N>5 for each image.

Our work highlights the profound compartment-specific spatiotemporally dynamics of MTs and the ER during specialized cell death and identifies a stereotyped degenerative hub where dramatic subcellular events take place to eliminate a polarized cell compartment. We propose (**FIG 6U**) that the ER executes a local two-step killing function by regulating microtubules both constructively (for recruitment and retention) as non-canonical ncMTOC and destructively (promoting severing). Both functions require ATLN-1/Atlastin, which stabilizes the ER and ensuring ER structural integrity. ATLN-1 also helps position SPAS-1 so that it can engage with and sever MTs of the distal node in an organized manner. We propose that ER-associated SPAS-1/Spastin of the distal node serves in bulk yet organized severing of MTs that accumulate at the distal node following retraction allowing for the complete dissolution of the distal node. We speculate that the ER-independent SPAS-1 of the distal tip facilitates the distal process retraction step in a manner dependent on an as yet uncharacterized role of ATLN-1/Atlastin that may be independent of both the ER and Golgi.

## Discussion

We identify the distal node as a stereotyped degenerative hub, where defined profound subcellular events take place. We present the ER as “killer organelle” in this new degenerative compartment and propose it plays dual roles to stabilize MTs for dismantling, both involving ATLN-1/Atlastin.

Previous work in Drosophila has linked Atlastin with microtubule stability in the context of neuromuscular junction growth and development(38). Our study can be distinguished from this work in multiple ways. While our super-resolution imaging studies also show that loss of ATLN-1/Atlastin causes aberrant ER morphology, we show cell-autonomous function of ATLN-1/Atlastin in the dying cell, and therefore in a different context from synaptic and muscle growth and scaffolding and the previous report of cell non-autonomous role of Atlastin. We define specific spatial and temporal roles for ATLN-1/Atlastin in a regressive remodeling event where ATLN-1 assists both in MT organization and severing.

We introduce here the ER as a possible ncMTOC. The ER has been shown to influence MT distribution(20). In Drosophila embryos, it has been shown that ER integrity influences mitotic spindle size(39). Our general knowledge of ncMTOCs in neurites has been rapidly growing from of genetic model organisms such and *C. elegans* and Drosophila(22). Our study offers important insights on a region-specific role in a differentiated cell in the nematode with a defined cell fate. Our study introduces the ER as a new ncMTOC promoter in a spatiotemporal specific way. It also shows that different types of ncMTOCs can arise in different compartments at different developmental timepoints to serve different functions. Studying the various forms of ncMTOCs in different contexts in the TSC will further our understanding of these fascinating subcellular structures. In addition, it has been reported that the ER and MTs can also stabilize each other(40).

Another intriguing observation in our study is the spatially-restricted nature of the ER in the TSC process. What leaves an ER-excluded zone distal for an organelle known to be vast in its expanse throughout the cell? ER is known to be excluded from the mitotic spindle to varying degrees in various cell types. It has been shown in HeLa cells that the ER is excluded from the mitotic spindle when the ER Ca^2+^ sensor, Stromal interaction molecule 1 (STIM1) is phosphorylated and inhibited(41). STIM proteins can sense when ER Ca^2+^ levels fall and directly activate Orai PM Ca^2+^ channels(42). It is possible Ca^2+^ dynamics play a role in the ER distribution in the TSC process.

We also suggest that SPAS-1 has both an ER dependent and independent role in CCE. SPAS-1 was identified in worms to be a homolog for human Spastin (43) and the specific importance for its C terminal domain (44) and pore forming regions (45) in its severing role have been described. In our study, while we do show a correlation between ER shape and SPAS-1/Spastin function, we do not directly show incorporation into the ER membrane, as known to be the case in human Spastin a hairpin protein(46). We propose that the association with the ER confers an enhanced severase capability on SPAS-1/Spastin, speculating that in *atln-1* and *lnp-1* mutants, where the association between the ER and SPAS-1/Spastin is lost. The exact functional relationship between the ER and Spastin is unknown. It is possible that the loss of ER tubules, reduced ER 3-way junctions such that SPAS-1/Spastin cannot perform its MT severing function when as it would when localized to these sites via ATLN-1/Atlastin. An outstanding question is what non-ER organelle SPAS-1/Spastin is associated with distally, if any.

How are MTs transported to the distal node? MTs can also interact with other components of the cytoskeleton, such as actin, which also plays an important role in establishing and maintaining neuronal polarity(47). The actin cytoskeleton is known to play roles in *C. elegans* early embryonic development(17). ER and actin structure and dynamics are also shown to be linked(48, 49). Actin has also been shown to transport MTs(50, 51). The actin network also influences the MTOC of centrosomes(49). The actin cytoskeleton may act as a roadway to transport MTs from the relevant compartment into the growing MTOC.

In neurons, region-specific destruction in the form of pruning can occur both in the form of neurite beading and fragmentation and in the form of retraction-like events wherein axons draw back and do not fragment. An example of this retraction is rodent hippocampal infrapyramidal tract (IPT) during postnatal development(52). We speculate that the distinct modes of elimination of the TSC proximal and distal process is linked to structures and function of these compartments. Our hypothesis is that the differential strategies were adopted to accommodate for the structure of the cell. The proximal process is thinner and is removed rapidly following simple beading and fragmentation. However, the distal process forms the actual tail-spike early on and acts as a scaffold; the distal node is formed from reorganization that allows the tip of the TSC process to remain as long as possible. Both the early TSC spike and the later distal node appear to involve different types of ncMTOCs. We propose that since the distal process is a sturdier structure with more MTs such that the beading-fragmentation phagocytosis strategy would not suffice for efficient removal. In order for the distal region to be phagocytosed, it needs to be reduced. As such, MTs travel to the distal node in an ER-independent manner, where they suffer bulk destruction by ER-associated SPAS-1.

Several intriguing questions to be expanded upon stem from the present study and motivate future studies. How does the ER confer enhanced severing capacity to SPAS-1? It may be through tethering and localization via ATLN-1. How are MTs specifically directed into the MTOC? What is the composition and role of the early MTOC of the TSC? Is it needed for the TSC’s scaffolding function? What is the role of other MT severases? Future forward and reverse genetic screens hold promise in uncovering answers.

We demonstrate the importance of the micro-organization of a cell and the resulting diverse interactions between subcellular components and offer descriptions for roles of an MTOC, SPAS-1/Spastin and the ER in this new context. Given considerable interest and gaps in knowledge on the roles of MT dismantling in pruning and neurodegeneration, and the ER in polarized cels such as neurons, our work has important implications for developmental pruning and pathological neurodegeneration. We also link conserved genes linked to neurodegenerative disease (HSP) in a developmental cell death process which opens the door to the use of CCE to mechanically understand both developmental and pathological cell death.

## Materials and Methods

### *C. elegans* methods

*C. elegans* strains were cultured using standard methods on *E. coli* OP50 and grown at 20ºC. Wild-type animals were the Bristol N2 subspecies. For scoring L1 TSC experiments, one of two integrated reporters were used: *nsIs435* or *nsIs685*. Integration of extrachromosomal arrays was performed using UV and trioxsalen (T2137, Sigma). Animals were scored at 20ºC.

### Microscopy and image processing

Images were collected on a Nikon TI2-E Inverted microscope using a CFI60 Plan Apochromat Lambda 60x Oil Immersion Objective Lens, N.A. 1.4 (Nikon) and a Yokogawa W1 Dual Cam Spinning Disk Confocal. Images were acquired using NIS-Elements Advanced Research Package. For still embryo imaging, embryos were anesthetized using 0.5 M sodium azide. Larvae were paralyzed with 10mM sodium azide. Super-resolution images were taken using a VTiSIM Super resolution Live Cell Confocal Imaging System.

### Quantifications

For CCE defects, TSC death defects were scored at the L1 stage. Animals were mounted on slides on 2% agarose-M9 pads, paralyzed with 10mM sodium azide, and examined on a Zeiss Axio-Scope A1. The persisting TSC was identified by fluorescence based on its location and morphology in the tail.

### Worm strains used in this study

The list of mutant *C*.□*elegans* strains used in this study is as follows:

LGI- *yop-1(ok3629), tba-1(ok1135)*

LGIV- *atln-1(ok1144), ced-3(n717)*

LGV- *spas-1(tm683), ret-1(gk242), aman-2(syb10402)*,

LGX- *lnp-1(tm1247)*

### Germline transformation and rescue experiments

Germline transformation was carried out as previously described<u>^37^</u>. All plasmids were injected at between 1 and 20□ng per *µ*l. pBSK was used to adjust the DNA concentration of injection mixtures if necessary. All rescue experiments were done with *myo-2*p::GFP as a co-injection marker along with *cdh-3*p::mCherry to label the TSC.

### Transgenes

The full list of transgenes is described in **Supplementary Table 1**. The full length or fragment of the *aff-1* promoter was used to label the TSC.

### Primers and plasmid construction

Primer sequences and information on the construction of plasmids used in this study are provided in **Supplementary Table 2**.

### Scoring of TSC/CCE defects

TSC death was scored at the L1 stage. Animals were synchronized allowing gravid hermaphrodites to lay eggs, washing off larvae and waiting 4-8 hours for hatched L1s. L1s synchronized were then mounted on slides on 2% agarose-water pads, anaesthetized in 10□mM sodium azide and examined on a Zeiss Axioplan 2 or Axio-Scope A1 under Nomarski optics and wide-field fluorescence at 40x or 60x. The TSC was identified by green fluorescence (from reporter transgenes) as well as by its location and morphology.

### Mutagenesis and mutant identification

*nsIs435* animals were mutagenized using 75□mM ethylmethanesulfonate (M0880, Sigma) for 4□h at 20□°C. Approximately 27,000 F2 progeny were screened for TSC persistence on a Zeiss Axio-Scope A1 at 40x. *lnp-1 (ns834)* was identified from analysis of Whole Genome Sequencing data, fosmid rescue and candidate gene analysis. See Supplementary Table 2.

### CRISPR-Cas9 genome editing

The strain PHX10402 *aman-2(syb10402)* was generated by SUNY Biotech.

### Image quantifications

Intensity profiles for the TSC soma distal degrading (SDD) stage were quantified and plotted using the Fiji/Image J Plugin tool.

### Statistics and reproducibility

The sample sizes and statistical tests were selected based on previous studies with similar methodologies (REF). Sample sizes were not determined using statistical methods and all experiments were repeated at least two to three times, as indicated, giving similar results. Independent transgenic lines were treated as independent experiments. Quantification of TSC persistence was done using an unpaired two-tailed *t*-test (Graphpad). For all figures, mean□±□standard error of the mean (s.e.m.) is represented.

## Author Contributions and Acknowledgments

KJ and PG designed the experiments and wrote the paper. KJ, HS and MJ performed experiments. We thank Ginger Clark, Rashna Sharmin and Aladin Elkhalil for technical assistance. We thank members of the Ghose lab for comments on the manuscript. We thank SUNY Biotech for strains. Some strains were provided by the CGC, which is funded by NIH Office of Research Infrastructure Programs (P40 OD010440).

## Supplemental Tables

**Supplemental table 1** Plasmid

**Supplemental table 2** Transgenes and strains

## Supplemental Movies

**Supplemental movie M1** Co-labeling TBA-1 and SPAS-1 in soma distal degrading (SDD) in wildtype embryos during CCE. Movies show planes across the TSC.

**Supplemental movie M2** Co-labeling TBA-1 and SPAS-1 in soma distal degrading (SDD) in *atln-1(ok1144)* embryos during CCE. Movies show planes across the TSC.

## Supplemental Figures

**Supplemental Figure S1** *lnp-1* gene structure showing the allele location and double mutant graph.

**Supplemental Figure S2** Co-labeling TBA-1 and SPAS-1 in soma distal degrading (SDD) in wildtype and *atln-1(ok1144)* to show distal tip GFP signal. TBA-1 signal in wildtype distal tip appears smaller and discrete. Where in *atln-1(ok1144)* it appears larger and less distributed.

## References

1. Donato A, Kagias K, Zhang Y, Hilliard MA. Neuronal sub-compartmentalization: a strategy to optimize neuronal function. Biol Rev Camb Philos Soc. 2019;94(3):1023–37.

2. Schuldiner O, Yaron A. Mechanisms of developmental neurite pruning. Cell Mol Life Sci. 2015;72(1):101–19.

3. Maor-Nof M, Yaron A. Neurite pruning and neuronal cell death: spatial regulation of shared destruction programs. Curr Opin Neurobiol. 2013;23(6):990–6.

4. Faust TE, Gunner G, Schafer DP. Mechanisms governing activity-dependent synaptic pruning in the developing mammalian CNS. Nat Rev Neurosci. 2021;22(11):657–73.

5. Neniskyte U, Gross CT. Errant gardeners: glial-cell-dependent synaptic pruning and neurodevelopmental disorders. Nat Rev Neurosci. 2017;18(11):658–70.

6. Yaron A, Schuldiner O. Common and Divergent Mechanisms in Developmental Neuronal Remodeling and Dying Back Neurodegeneration. Curr Biol. 2016;26(13):R628–R39.

7. Kramer R, Wolterhoff N, Galic M, Rumpf S. Developmental pruning of sensory neurites by mechanical tearing in Drosophila. J Cell Biol. 2023;222(3).

8. Furusawa K, Emoto K. Spatiotemporal regulation of developmental neurite pruning: Molecular and cellular insights from Drosophila models. Neurosci Res. 2021;167:54–63.

9. Raiders S, Black EC, Bae A, MacFarlane S, Klein M, Shaham S, et al. Glia actively sculpt sensory neurons by controlled phagocytosis to tune animal behavior. Elife. 2021;10.

10. Gurel PS, Hatch AL, Higgs HN. Connecting the cytoskeleton to the endoplasmic reticulum and Golgi. Curr Biol. 2014;24(14):R660–R72.

11. Yalcin B, Zhao L, Stofanko M, O’Sullivan NC, Kang ZH, Roost A, et al. Modeling of axonal endoplasmic reticulum network by spastic paraplegia proteins. Elife. 2017;6.

12. Kuijpers M, Nguyen PT, Haucke V. The Endoplasmic Reticulum and Its Contacts: Emerging Roles in Axon Development, Neurotransmission, and Degeneration. Neuroscientist. 2023:10738584231162810.

13. Liu X, Guo X, Niu L, Li X, Sun F, Hu J, et al. Atlastin-1 regulates morphology and function of endoplasmic reticulum in dendrites. Nat Commun. 2019;10(1):568.

14. Puhka M, Vihinen H, Joensuu M, Jokitalo E. Endoplasmic reticulum remains continuous and undergoes sheet-to-tubule transformation during cell division in mammalian cells. J Cell Biol. 2007;179(5):895–909.

15. Summerville JB, Faust JF, Fan E, Pendin D, Daga A, Formella J, et al. The effects of ER morphology on synaptic structure and function in Drosophila melanogaster. J Cell Sci. 2016;129(8):1635–48.

16. Rumpf S, Wolterhoff N, Herzmann S. Functions of Microtubule Disassembly during Neurite Pruning. Trends Cell Biol. 2019;29(4):291–7.

17. Velarde N, Gunsalus KC, Piano F. Diverse roles of actin in C. elegans early embryogenesis. BMC Dev Biol. 2007;7:142.

18. Datar A, Ameeramja J, Bhat A, Srivastava R, Mishra A, Bernal R, et al. The Roles of Microtubules and Membrane Tension in Axonal Beading, Retraction, and Atrophy. Biophys J. 2019;117(5):880–91.

19. Erturk A, Hellal F, Enes J, Bradke F. Disorganized microtubules underlie the formation of retraction bulbs and the failure of axonal regeneration. J Neurosci. 2007;27(34):9169–80.

20. Tikhomirova MS, Kadosh A, Saukko-Paavola AJ, Shemesh T, Klemm RW. A role for endoplasmic reticulum dynamics in the cellular distribution of microtubules. Proc Natl Acad Sci U S A. 2022;119(15):e2104309119.

21. Sanchez AD, Branon TC, Cote LE, Papagiannakis A, Liang X, Pickett MA, et al. Proximity labeling reveals non-centrosomal microtubule-organizing center components required for microtubule growth and localization. Curr Biol. 2021;31(16):3586–600 e11.

22. Sanchez AD, Feldman JL. Microtubule-organizing centers: from the centrosome to non-centrosomal sites. Curr Opin Cell Biol. 2017;44:93–101.

23. Vineethakumari C, Luders J. Microtubule Anchoring: Attaching Dynamic Polymers to Cellular Structures. Front Cell Dev Biol. 2022;10:867870.

24. Zhu X, Kaverina I. Golgi as an MTOC: making microtubules for its own good. Histochem Cell Biol. 2013;140(3):361–7.

25. Kuo YW, Howard J. Cutting, Amplifying, and Aligning Microtubules with Severing Enzymes. Trends Cell Biol. 2021;31(1):50–61.

26. McNally FJ, Roll-Mecak A. Microtubule-severing enzymes: From cellular functions to molecular mechanism. J Cell Biol. 2018;217(12):4057–69.

27. Errico A, Ballabio A, Rugarli EI. Spastin, the protein mutated in autosomal dominant hereditary spastic paraplegia, is involved in microtubule dynamics. Hum Mol Genet. 2002;11(2):153–63.

28. Liu Q, Zhang G, Ji Z, Lin H. Molecular and cellular mechanisms of spastin in neural development and disease (Review). Int J Mol Med. 2021;48(6).

29. Roll-Mecak A, Vale RD. Structural basis of microtubule severing by the hereditary spastic paraplegia protein spastin. Nature. 2008;451(7176):363–7.

30. Kuo YW, Trottier O, Mahamdeh M, Howard J. Spastin is a dual-function enzyme that severs microtubules and promotes their regrowth to increase the number and mass of microtubules. Proc Natl Acad Sci U S A. 2019;116(12):5533–41.

31. Aiken J, Holzbaur ELF. Spastin locally amplifies microtubule dynamics to pattern the axon for presynaptic cargo delivery. Curr Biol. 2024;34(8):1687–704 e8.

32. Costa AC, Sousa MM. The Role of Spastin in Axon Biology. Front Cell Dev Biol. 2022;10:934522.

33. Sanderson CM, Connell JW, Edwards TL, Bright NA, Duley S, Thompson A, et al. Spastin and atlastin, two proteins mutated in autosomal-dominant hereditary spastic paraplegia, are binding partners. Hum Mol Genet. 2006;15(2):307–18.

34. Ghose P, Rashid A, Insley P, Trivedi M, Shah P, Singhal A, et al. EFF-1 fusogen promotes phagosome sealing during cell process clearance in Caenorhabditis elegans. Nat Cell Biol. 2018;20(4):393–9.

35. Chen S, Desai T, McNew JA, Gerard P, Novick PJ, Ferro-Novick S. Lunapark stabilizes nascent three-way junctions in the endoplasmic reticulum. Proc Natl Acad Sci U S A. 2015;112(2):418–23.

36. Sallee MD, Zonka JC, Skokan TD, Raftrey BC, Feldman JL. Tissue-specific degradation of essential centrosome components reveals distinct microtubule populations at microtubule organizing centers. PLoS Biol. 2018;16(8):e2005189.

37. Chabin-Brion K, Marceiller J, Perez F, Settegrana C, Drechou A, Durand G, et al. The Golgi complex is a microtubule-organizing organelle. Mol Biol Cell. 2001;12(7):2047–60.

38. Lee M, Paik SK, Lee MJ, Kim YJ, Kim S, Nahm M, et al. Drosophila Atlastin regulates the stability of muscle microtubules and is required for synapse development. Dev Biol. 2009;330(2):250–62.

39. Araujo M, Tavares A, Vieira DV, Telley IA, Oliveira RA. Endoplasmic reticulum membranes are continuously required to maintain mitotic spindle size and forces. Life Sci Alliance. 2023;6(1).

40. Farias GG, Freal A, Tortosa E, Stucchi R, Pan X, Portegies S, et al. Feedback-Driven Mechanisms between Microtubules and the Endoplasmic Reticulum Instruct Neuronal Polarity. Neuron. 2019;102(1):184–201 e8.

41. Smyth JT, Beg AM, Wu S, Putney JW, Jr., Rusan NM. Phosphoregulation of STIM1 leads to exclusion of the endoplasmic reticulum from the mitotic spindle. Curr Biol. 2012;22(16):1487–93.

42. Carrasco S, Meyer T. STIM proteins and the endoplasmic reticulum-plasma membrane junctions. Annu Rev Biochem. 2011;80:973–1000.

43. Matsushita-Ishiodori Y, Yamanaka K, Ogura T. The C. elegans homologue of the spastic paraplegia protein, spastin, disassembles microtubules. Biochem Biophys Res Commun. 2007;359(1):157–62.

44. Onitake A, Matsushita-Ishiodori Y, Johjima A, Esaki M, Ogura T, Yamanaka K. The C-terminal alpha-helix of SPAS-1, a Caenorhabditis elegans spastin homologue, is crucial for microtubule severing. J Struct Biol. 2012;179(2):138–42.

45. Matsushita-Ishiodori Y, Yamanaka K, Hashimoto H, Esaki M, Ogura T. Conserved aromatic and basic amino acid residues in the pore region of Caenorhabditis elegans spastin play critical roles in microtubule severing. Genes Cells. 2009;14(8):925–40.

46. Connell JW, Lindon C, Luzio JP, Reid E. Spastin couples microtubule severing to membrane traffic in completion of cytokinesis and secretion. Traffic. 2009;10(1):42–56.

47. Konietzny A, Bar J, Mikhaylova M. Dendritic Actin Cytoskeleton: Structure, Functions, and Regulations. Front Cell Neurosci. 2017;11:147.

48. Pain C, Tolmie F, Wojcik S, Wang P, Kriechbaumer V. intER-ACTINg: The structure and dynamics of ER and actin are interlinked. J Microsc. 2023;291(1):105–18.

49. Yamamoto S, Gaillard J, Vianay B, Guerin C, Orhant-Prioux M, Blanchoin L, et al. Actin network architecture can ensure robust centering or sensitive decentering of the centrosome. EMBO J. 2022;41(20):e111631.

50. Etienne-Manneville S. Actin and microtubules in cell motility: which one is in control? Traffic. 2004;5(7):470–7.

51. Hasaka TP, Myers KA, Baas PW. Role of actin filaments in the axonal transport of microtubules. J Neurosci. 2004;24(50):11291–301.

52. Bagri A, Cheng HJ, Yaron A, Pleasure SJ, Tessier-Lavigne M. Stereotyped pruning of long hippocampal axon branches triggered by retraction inducers of the semaphorin family. Cell. 2003;113(3):285–99.

