## Supplementary figures and images for "The endoplasmic reticulum promotes microtubule organization and region-specific disassembly to execute Compartmentalized Cell Elimination"

### Supplemental Figure 1

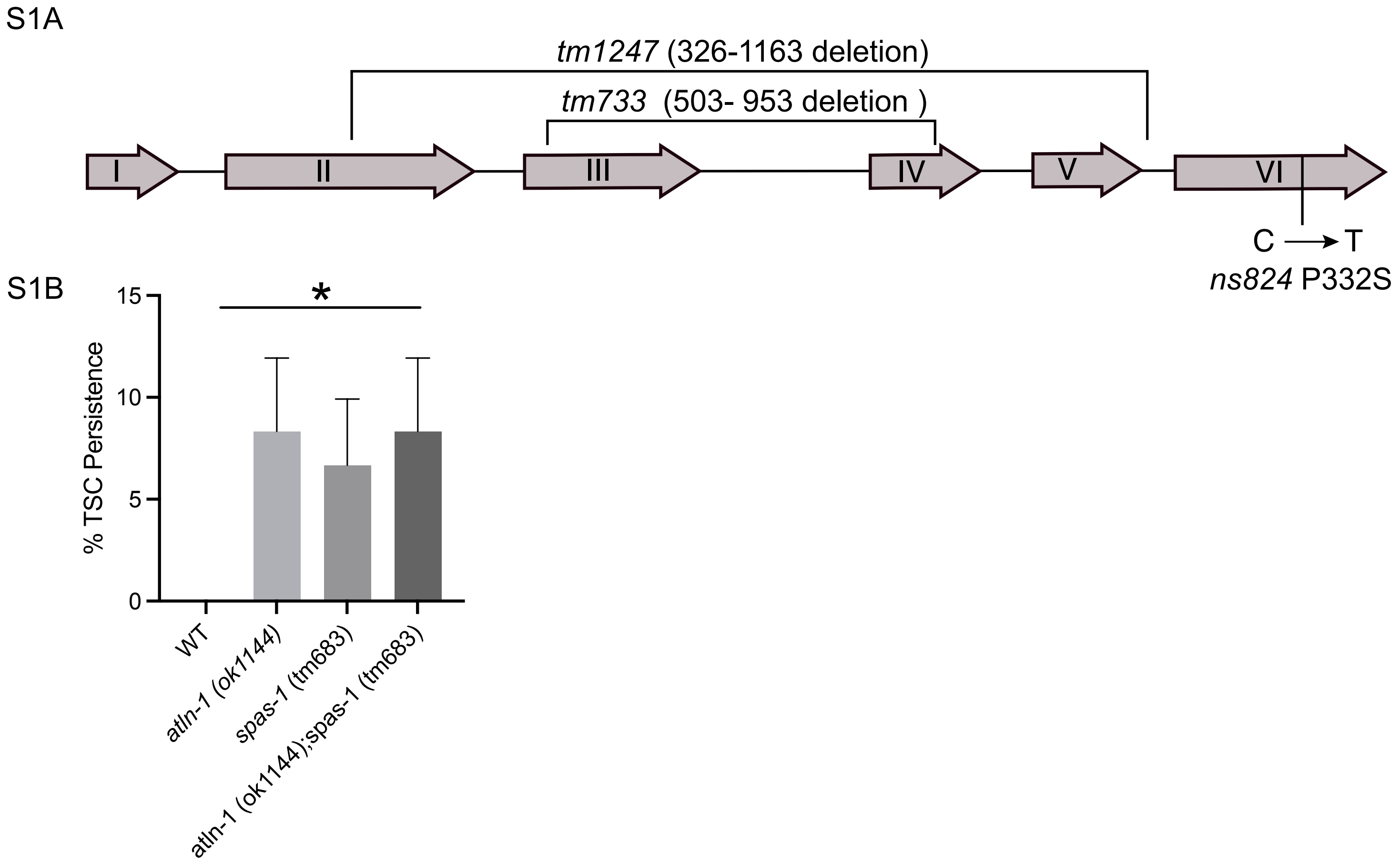

### Supplemental Figure 2

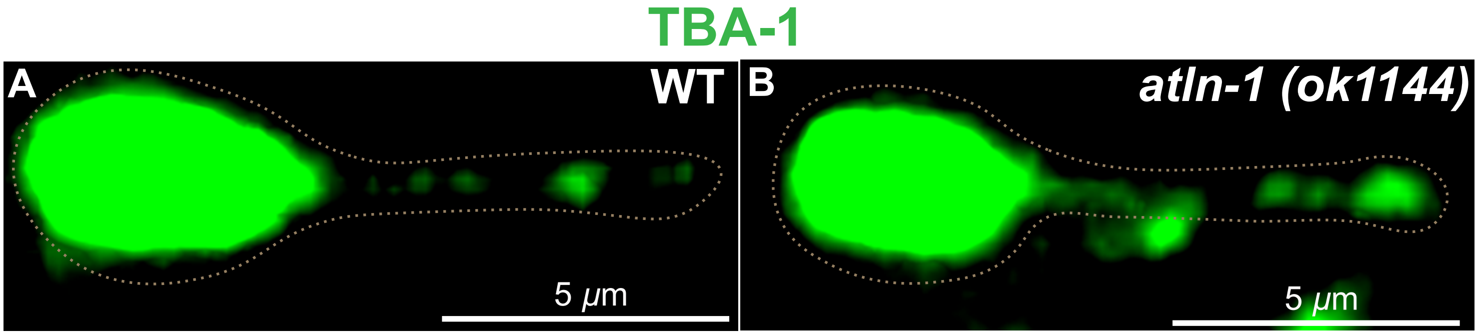

### Supplemental Movie 1

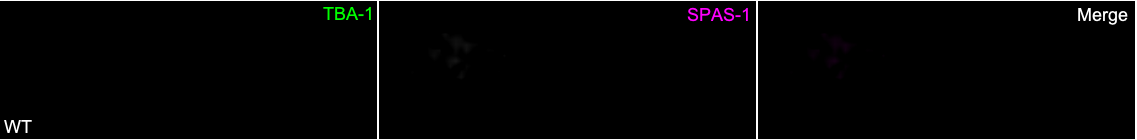

### Supplemental Table 2

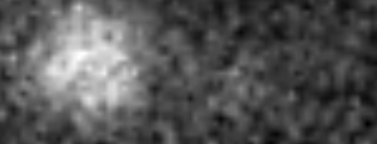
