## Supplemental Table 1 for "The endoplasmic reticulum promotes microtubule organization and region-specific disassembly to execute Compartmentalized Cell Elimination"

| **Plasmid number (pPG)** | **Primers used** | **Sequences** |
| --- | --- | --- |
|  | Backbone |  |
| pPG269 (*lnp-1pro*::mKate2) | C418 | ATGGTCTCCGAGCTCATTAAcGAAAAC |
|  | C419 | GATCCTCTAGAGTCGACCTGCAGGC |
|  | oKJ36 | **GAAATAAGCTTGCATGCCTGCAGGTCGACTCTAGAGGATCtgccatgttttatttctattctcaattcc** |
|  | oKJ37 | **GCTTCATATGCATGTTTTCgTTAATGAGCTCGGAGACCATctgaaaaataaattacattttaattttaaaaaaag** |
| pPG270 (TSCpro:: *lnp-1* cDNA_SL2::mCherry) | Backbone |  |
|  | C422 | GGCGCGCCgttccgaatattatg |
|  | C423 | GAATTCcaactgagcgccggtc |
|  | Insert |  |
|  | oKJ38 | **tccatactttctcatttcataatattcggaacGGCGCGCCaaaATGGGTAACTTATTTTCAAGGAACAA** |
|  | oKJ39 | **tacctttgggtcctttggccaatcccggggatcctctagaTTAATGGAATTCAGTTTCCATTGTCTTGG** |
| pPG277 (TSCpro::vits22_KDEL oxGFP) (ER) | Backbone |  |
|  | oKJ43 | GGCGCGCCgttccgaatattatg |
|  | oKJ44 | GAATTCcaactgagcgccggtc |
|  | Insert |  |
|  | oKJ67 | **tccatactttctcatttcataatattcggaacGGCGCGCCAAAAAATGCGTTCAATCATAATCGCCTC** |
|  | oKJ68 | **caagttggtaatggtagcgaccggcgctcagttgGAATTCTTAGAGCTCGTCCTTGTCGTCGGATC** |
| pPG287 (*atln-1*pro::mKate2) | Backbone |  |
|  | C418 | ATGGTCTCCGAGCTCATTAAcGAAAAC |
|  | C419 | GATCCTCTAGAGTCGACCTGCAGGC |
|  | Insert |  |
|  | oKJ75 | GAAATAAGCTTGCATGCCTGCAGGTCGACTCTAGAGGATCttttttggtgtaattaagcaaacaatcgg |
|  | oKJ76 | GCTTCATATGCATGTTTTCgTTAATGAGCTCGGAGACCATttttttttgctgaaaaaaggggtgaaatttt |
| pPG291 (*spas-1pro* ::mKate2) | Backbone |  |
|  | C418 | ATGGTCTCCGAGCTCATTAAcGAAAAC |
|  | C419 | GATCCTCTAGAGTCGACCTGCAGGC |
|  | Insert |  |
|  | oKJ81 | AGCTTGCATGCCTGCAGGTCGACTCTAGAGGATCCCCGGGcaatgtcagccgaattactcacaatg |
|  | oKJ82 | GCTTCATATGCATGTTTTCgTTAATGAGCTCGGAGACCATtggaactgaaaatttaatacaattggaa |
| pPG293 (TSCpro::GFP::TBA-1) | Backbone |  |
|  | oKJ43 | GGCGCGCCgttccgaatattatg |
|  | oKJ44 | GAATTCcaactgagcgccggtc |
|  | Inserts |  |
|  | oKJ83 | tccatactttctcatttcataatattcggaacGGCGCGCCATGAGTAAAGGAGAAGAACTTTTCACTG |
|  | oKJ84 | tttacaacaagtaaataaaaacaaaaccaggaacttacATTTTGTATAGTTCATCCATGCCATGTGTAA |
|  | oKJ85 | AGCTGCTGGGATTACACATGGCATGGATGAACTATACAAAATgtaagttcctggttttgtttttatttact |
|  | oKJ86 | caagttggtaatggtagcgaccggcgctcagttgGAATTCTTAATACTCTTCTCCTTCCTCCTCGTTTC |
| pPG373 (TSCpro::mCherry SPAS-1) | Backbone |  |
|  | oKJ150 | GAATTCcaactgagcgccggtcgc |
|  | oKJ151 | CGATGCTCCTGAGGCTCCCGATGC |
|  | Insert |  |
|  | oKJ157 | cgagctgtacaagGGAGCATCGGGAGCCTCAGGAGCATCGATGTTCGCCTTTTCAAAAGGTCCCGCC |
|  | oKJ158 | caagttggtaatggtagcgaccggcgctcagttgGAATTCTTAGCAACCGAAACTTCGAGAGAAATCGGAG |
| pPG447 (TSCpro::GIP GFP SL2::myrmCherry TSC ) | Backbone |  |
|  | oKJ217 | CGATGCTCCTGAGGCTCCCGATGCTC |
|  | oKJ218 | GAATTCcaactgagcgccggtcgctac |
|  | Insert |  |
|  | oKJ219 | cgagctgtacaagGGAGCATCGGGAGCCTCAGGAGCATCGATGCGTCGACAAGGCAGCGAAGAAGTT |
|  | oKJ220 | caagttggtaatggtagcgaccggcgctcagttgGAATTCTCAAAAATTATAGAAGATTCGAATTTTCACCAA |
| pPG286 (TSCpro::*atln-1* cDNA_SL2::mCherry) | Backbone |  |
|  | C422 | GGCGCGCCgttccgaatattatg |
|  | C423 | GAATTCcaactgagcgccggtc |
|  | Insert |  |
|  | oKJ73 | tccatactttctcatttcataatattcggaacGGCGCGCCaaaATGGAAACAACTCCTCAAAACGAGC |
|  | oKJ74 | tacctttgggtcctttggccaatcccggggatcctctagaCTAATGCCGTTTTCTGAGTCCATC |
| pPG446 (TSCpro::*spas-1* cDNA_SL2::mCherry) | Backbone |  |
|  | C422 | GGCGCGCCgttccgaatattatg |
|  | C423 | GAATTCcaactgagcgccggtc |
|  | Insert |  |
|  | oKJ212 | tccatactttctcatttcataatattcggaacGGCGCGCCaaaATGTTCGCCTTTTCAAAAGGTCCCG |
|  | oKJ213 | tacctttgggtcctttggccaatcccggggatcctctagaTTAGCAACCGAAACTTCGAGAGAAATCGG |
